# Dietary Vitamin A intakes among Pregnant Women Attending Antenatal Care in Health Facilities: As a connotation for the progress of vitamin A Deficiency in Dessie Town, North East Ethiopia

**DOI:** 10.1101/516047

**Authors:** Zenebech Koricho, Gudina Egata Atomssa, Tefera Chane Mekonnen, Sisay Eshete Tadesse

**Affiliations:** Lecturer at Dessie College of Health Science, Dessie, Northern Ethiopia.; Associate Professor of Public Health at College of Health sciences and Medicine, Haramaya University, Harare, Eastern Ethiopia.; Lecturer and Researcher at Department of Public Health, College of Medicine and Health Sciences, Wollo University, Dessie, Northern Ethiopia.

**Keywords:** Adequacy of Vitamin A, Pregnant women, Dessie, Ethiopia

## Abstract

Vitamin A plays important roles in vision, cellular differentiation, embryonic development, reproduction, growth, and the immune system. Women living in developing countries are at increased risk of undernutrition during pregnancy due to poverty, poor diet quality and quantity, and high fertility rate. Dietary quality and diversity reflect adequacy of vitamin A whereby reduce the risk of vitamin A deficiency. The aim of study was to determine adequacy of vitamin A among pregnant women attending antenatal care in health facilities of Dessie town, Ethiopia, January, 2017. Health facility-based cross-sectional study was conducted among 390 women that attended antenatal care in Dessie town. The 9 food groups from FAO based on 24hours dietary recall was used for data collection. Adequacy of vitamin A was determined from nutrient adequacy ratio after obtaining report of nutrient intake from food composition table version III and IV in terms of B carotene and retinol equivalent respectively, based on estimated average requirement recommendation of vitamin A, 370 RE/day for pregnant women. Multivariable Logistic regression analysis was done after dichotomizing the dependent variables.

Adequacy of vitamin A among pregnant women was 41.8 % with an average nutrient adequacy ratio of 0.9. The mean dietary intake of vitamin A was 290.1μg per day. The predictors for adequacy of vitamin A were high and medium women diversity score (AOR=**2.92**; CI: 1.50-5.70) and (AOR=**1.87**; CI: 1.11- 3.16).

In this study adequacy of vitamin A was low and affected by women dietary diversity score. Focusing on food based approaches especially educating pregnant women to diversify their diet is crucial to reduce their risk of vitamin a deficiency.

## Background

Vitamin A is an essential nutrient that plays roles in the vision, cellular differentiation, embryonic development, reproduction, growth and the immune system (1–2). Its deficiency called Vitamin A deficiency (VAD) is the third most common nutritional deficiency in the world and mostly severe in Southeast Asia and Sub-Saharan Africa (3).

Dietary sources of vitamin A are preformed vitamin A; and pro-vitamin A, which are those Carotenoids precursors that can be bio-converted to retinol. Pro-vitamin A sources contribute 30–35% of the total vitamin A in the food supplies of industrialized populations, which is 70–75% in the populations of Africa and Southern Asia (4).

Nutritional adequacy is defined as the sufficient intake of essential nutrients including vitamin A. Adequacy of nutrients is needed to fulfill nutritional requirements for optimal health; the prevention of chronic diseases or the reduction of risk for diet associated diseases (5). Low vitamin A intake during nutritionally demanding periods in life, such as infancy, childhood, pregnancy and lactation increases the risk of health consequences, or vitamin A deficiency disorders (VADD) (6, 7). Dietary diversity scores can be used as a proxy measure of micronutrient and diet adequacy of women (8). Dietary diversity has been linked to less reporting of a major pregnancy related complications, like preeclampsia and Eclampsia (9, 10).

The most common cause of Vitamin A deficiency is low dietary intake of Vitamin A. When dietary intake is chronically low, there will be insufficient vitamin A to support vision and cellular processes, leading to impaired tissue function (11, 12). Studies have shown that improving vitamin A status can reduce pregnancy-related mortality by as much as 40 %(7, 13). Although considerable progress has been made in controlling Vitamin A deficiency worldwide, there is still a need for additional prevention efforts in the form of dietary approaches (13, 14).

In Ethiopia strategies involving multiple stakeholders and public–private partnership are well designed with National Nutrition Program (NNP) with implementing sectors have declared and pledged their commitment to support the achievement of the targets of the program (15).

However, these efforts are not yet put on the ground to the point of addressing the adequacy of vitamin A among pregnant women. Taking into consideration that the period of pregnancy is a window of exposure for nutritional detrimental effects; this study was conducted on adequacy of vitamin A among pregnant women that attended antenatal care in health facilities of Dessie town as these people are part and parcel of the vulnerable groups in Ethiopia.

## Methods

### Study Area and Period

A health facility-based cross-sectional survey was conducted from January 12 to February 12, 2017 in Dessie Town located 401 Kms from the capital city, Addis Ababa and 480 kms from the regional capital city, Bahir Dar, Ethiopia. Total population in Dessie town was 216,384, among which 7,292 were expected to be pregnant women. There are 12, 17 and 4 governmental, private and nongovernmental health facilities (total of 33 health facilities) in Dessie town respectively. Among the total pregnant women, 4397 pregnant women have got ANC in 2016 in the governmental, private and nongovernmental health facilities of the town.

Maternal and child health care is given in most of government, some private and nongovernmental health facilities like immunization, Family planning, Antenatal Care, Delivery care, Postnatal care. In Dessie Town available common food products include cereals, grains, legumes, dairy products, egg, fish, meat, fruits (Mango, papaya, Avocado, Orange) and vegetables (Spinach, Lettuce, Green pepper, carrot, pumpkin, cabbage).

All pregnant women who were attending ANC in 12 randomly selected health facilities (government, private and nongovernmental health facilities) of Dessie town were source population whereas randomly selected pregnant women who had been attending ANC in 12 governments, private and nongovernmental health facilities of Dessie town during data collection were study participants. All pregnant women who were attending in the randomly selected government, private and nongovernmental health facilities of Dessie town were included. Pregnant women found in fasting day and have dietary restriction were excluded from the study

### Sample size and Sampling Technique

The sample size was determined using single proportion population formula by considering the following assumptions of proportion 50% of adequacy of vitamin A, confidence level of 95%, margin of error 5% and 10% as non-response rate, the total sample size was **422**.

In Dessie town, 17 health facilities provide ANC service, among which 12 health facilities were randomly selected (07 governmental, 04 private and 01 non-governmental). Sample size was allocated proportionally to the client flow of each health facility. Those pregnant women who fulfilled the inclusion criteria and visited the ANC clinic during the study period were systematically selected as study participants after obtaining identification card of each pregnant woman.

## Data collection Techniques and Data Measurements

An Interviewer-administered pre-tested questionnaire was used for dietary data from known sources (7, 16–18). The questionnaire was translated to local language (Amharic) and then again back translated to English language. Eight data collectors and two supervisors were hired based on their prior exposure of research data collection.

The socio-demographic features (such as age, educational level, occupation, marital status, income level, family size), obstetrics related characteristics (birth interval, parity, months of pregnancy, birth interval) and dietary intake and habits of pregnant women (food taboos, nutrition education, daily intake of vitamin A rich sources, reasons for not daily taking of vitamin A rich vegetables, poultry production, home gardening of vegetables) were assessed to investigate the research objective.

Adequacy of vitamin A and Women dietary diversity score (WDDS) were measured by using the 9 food groups from FAO dietary diversity guideline based on interactive 24 hours dietary recall. Details of the 9 food groups was probed or interviewed from each participant. Dietary information was assembled and estimated using portion size and gram amounts to know the Carotenoids equivalent and retinol equivalent (RE). And then to determine the level of adequacy, individual RE from each food group was summed. Then the corresponding nutrient adequacy ratio (NAR) was calculated. As recommended by joint FAO/WHO consultation, mean requirement of vitamin A in mcg RE/day for pregnant women is 370 and safe intake in mcg is 600 RE/day. So that by using a cutoff value of 1 as adequate and <1 as inadequate; the outcome was defined by NAR (12). The WDDS was classified as low, medium, high according to FAO’s classification of WDDS of low WDDS(<3),medium WDDS(4–5) and high WDDS(≥6)(17).

Dietary intake of vitamin A of each study participant was converted to gram amounts then gram amounts calculated to Carotenoids equivalent as per 100 gram edible portion in the food composition table of Ethiopia.

Carotenoids equivalents were converted to retinol equivalent (RE) by a conversion factor of 6. Foods that have retinol values were simply recorded and summed to values obtained by conversion of Carotenoids to retinol (RE). To determine nutrient adequacy ratio (NAR) of vitamin A, ratio of observed intake was calculated in relation to recommended intake based on estimated average requirement (EAR) of 370 mcg RE/day (12).

In this study adequacy of vitamin A was defined as daily intake of the vitamin from plant and/or animal sources that is equal to the estimated average requirement of vitamin A, 370mcg RE/day. Adequacy of vitamin A was dichotomized as inadequate if NAR<1 and as adequate if NAR >=1(12, 17).

## Data quality control

Training was given for data collectors on the objectives of the study, the number and items of the data collection tools. Information about the essential technical skills on how to collect a 24 hours dietary recall method was also part of the training. Frequent supervision and checking of the data for consistency and completeness was done by supervisors and principal investigator. To reduce recall bias on dietary data, corresponding portion size of dietary intake was estimated by local measurement utensils by allowing study participants to choose from a variety of commonly used local measurement utensils offered during data collection and show which variety of local measurements was used for the portion size of their diet. Gram amounts for the portion sizes were determined as per the food composition table of Ethiopia (per 100 gram edible portion). Some consumed foods were estimated by their raw portion size or gram amounts.

## Data management and analysis

Data entry was done using Epi-Info 7.2 software program and data cleaning by SPSS version 20 software program. The food composition table of Ethiopia version III and IV was used to convert raw dietary data to Carotenoids equivalent and retinol equivalent of vitamin A rich foods consumed in the past 24 hours.

Descriptive statistics such as percentage, mean and standard deviation were done to express the findings. After dichotomizing the dependent variable, binary Logistic regression and multiple logistic regression analysis with crude odds ratio (COR) and adjusted odds ratio (AOR) were done to see the association of independent variables on the outcome variable and to control the possible confounding effects. Odds ratio was used to describe the study population in relation to relevant variables and to assess the presence and degree of association between dependent and independent variables. P-value less than 0.05 used to decide whether differences that occur would be statistically significant or not.

## Results

### Socio-demographic and economic characteristics of pregnant women

Out of a total of 422 pregnant women sampled, 390 women participated with 92.4% response rate. The mean age of the respondents was 26.89 (± 4.63 SD) years. Among the study participants 41(10.5%), 95(24.4%), 185(47.4%), and 69(17.7%) were unable to read and write, primary schooling, secondary schooling and college and above in their education respectively. The mean gestational age was 6.16(± 2.21SD) months (Table 1).

**Table 1:** Socio demographic and economic variables of pregnant women, health facilities of Dessie town, May 2017.

| Variables | Number of study participants<br>(n=390) | Percentage |
| --- | --- | --- |
| Maternal Education |  |  |
| Unable read and write | 41 | 10.5 |
| Primary schooling | 95 | 24.4 |
| Secondary schooling | 185 | 47.4 |
| College and above | 69 | 17.7 |
| Maternal Occupation |  |  |
| Unemployed | 14 | 3.6 |
| Government employed | 321 | 82.3 |
| Merchant | 55 | 14.1 |
| Stage of pregnancy(in Wks) |  |  |
| First trimester | 61 | 15.6 |
| Second trimester | 142 | 36.4 |
| Third trimester | 187 | 48.0 |
| Mean gestational age ± SD | 6.16 ± 2.21 months |  |
| Family size |  |  |
| <3 | 174 | 44.6 |
| 3-4 | 66 | 16.9 |
| >=5 | 150 | 38.5 |
| **Pregnancy related illness** |  |  |
| Yes | 201 | 51.5 |
| No | 189 | 48.5 |
| **Common illness during pregnancy** |  |  |
| **Dyspepsia** | 18 | 8.9 |
| **Morning sickness** | 166 | 82.6 |
| **Helminthes infestation** | 17 | 8.5 |
| **Birth interval (n=209)** |  |  |
| 1-5 years | 141 | 67.5 |
| >5years | 69 | 32.5 |
| **Parity** |  |  |
| <3 | 282 | 72.3 |
| >=3 | 108 | 27.7 |
| **Nutrition education** |  |  |
| **Yes** | 96 | 24.6 |
| **No** | 294 | 75.4 |

### Dietary diversity and Vitamin A adequacy of Pregnant Women

Poultry production & home gardening of vegetables for own consumption was assessed to see the availability of rich sources of Vitamin A. About one every five pregnant woman (22%) consumed vitamin A rich fruits and vegetables in daily basis (Table 2).

**Table 2:** Nutritional variables of pregnant women, health facilities of Dessie town, May 2017.

| Variables | Number of study participants |  |
| --- | --- | --- |
|  | (n=390) | % |
| **Daily intake of vitamin A rich vegetables and Fruits** |  |  |
| Yes | 86 | 22.0 |
| No | 304 | 77.9 |
| **Reasons for no Daily intake of vitamin A rich vegetables (n=304)** |  |  |
| Vitamin A rich sources available during market day | 67 | 17.2 |
| No fresh products except market days | 45 | 11.5 |
| Socio economic reasons | 192 | 49.2 |
| **Poultry production** |  |  |
| Yes | 56 | 14.4 |
| No | 334 | 85.6 |
| **Home garden** |  |  |
| Yes | 63 | 16.2 |
| No | 327 | 83.8 |

The foods eaten by the pregnant women were predominantly starchy staple foods (99.7%), other fruits and vegetables (98.7%), and legumes (84.9%) (Fig1). Women dietary diversity score(WDDS) from the nine food groups were measured and 26.9%, 55.4% and 17.7% of pregnant women found to have low, medium and high dietary diversity score respectively. The mean WDDS was 4.36 ± 1.2 SD.

**Figure 1:**
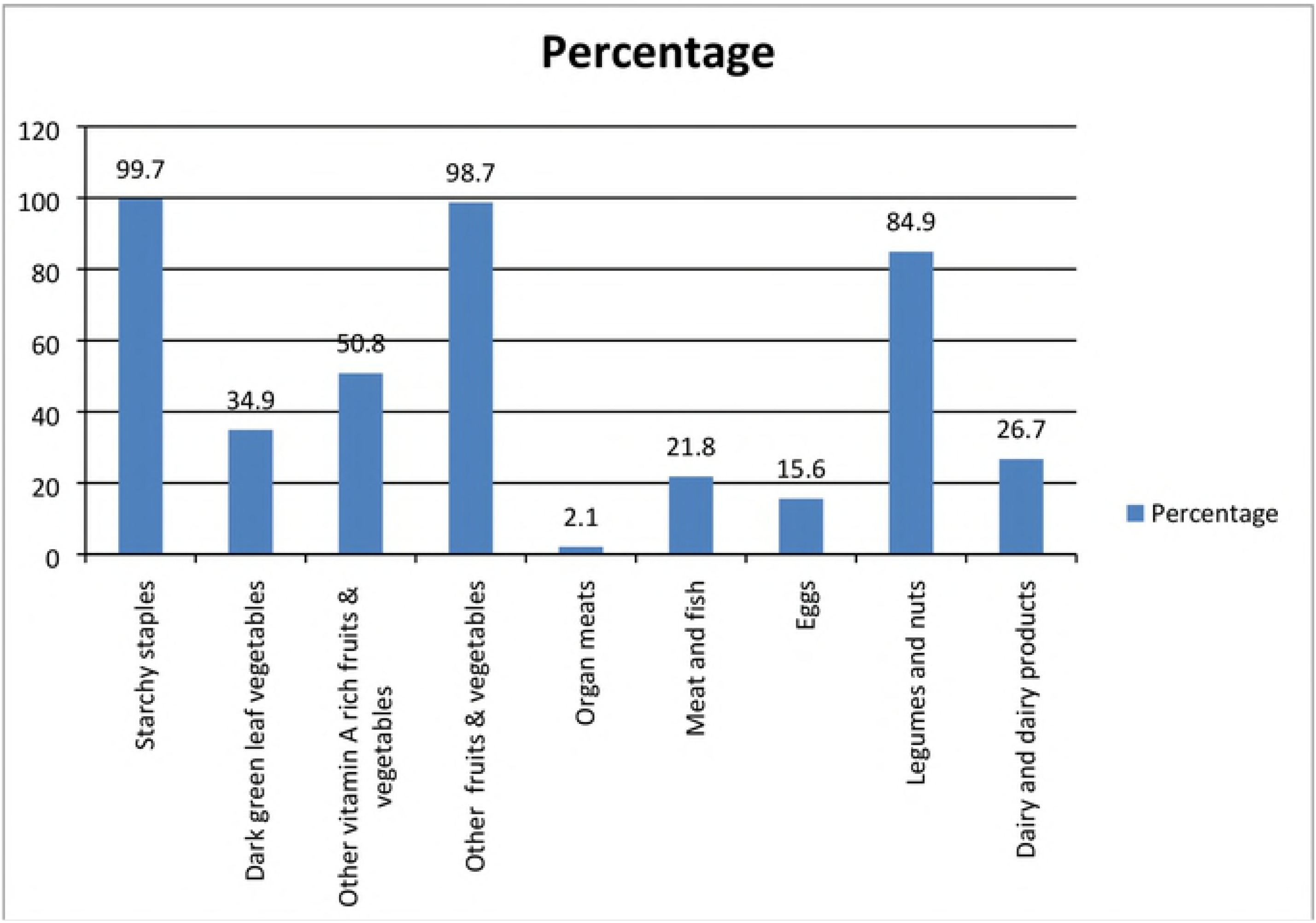
food groups consumed by pregnant women having ANC follow up in Health facilities of Dessie Town, Amhara Region, Ethiopia, May 2017.

The average dietary Vitamin A intakes were 294.1μg of retinol equivalents or 185.3 μg retinol activity equivalents. The average nutrient adequacy ratio of vitamin A from both preformed and pro-vitamin A sources was 0.9. About two every five pregnant women (41.8%) had adequate dietary intake of vitamin A (equivalent of recommended daily allowance of vitamin A). Regarding vitamin A intake from major food sources, Mango contributed RE of 19.5% of pregnant women in its juice or flesh form; this seems an emerging tendency to use it as food of choice by pregnant women.

### Factors Associated with Adequacy of Vitamin A

In the Bivariable logistic regression analysis, occupation, no poultry production for own consumption, gardening of vegetables for own consumption and women dietary diversity score (WDDS) were associated with adequacy of vitamin A.

Even though many independent variables reached the final step, with multivariable logistic regression analysis, WDDS and no poultry production were found to be significantly associated with adequacy of vitamin A.

The odds of vitamin A adequacy among pregnant women with high WDDS was **2.9** times higher than the odds of vitamin A adequacy among pregnant women with low WDDS (AOR=**2.92**; CI: 1.50-5.70). Similarly, the odds of vitamin A adequacy among pregnant women with medium WDDS was 1.87 times higher than the odds of vitamin A adequacy among pregnant women with low WDDS (AOR=**1.87**; CI: 1.11-3.16).

However, occupational status of pregnant women, rearing of poultry and having home gardening for cultivating vegetables were not significantly associated with adequacy of vitamin A intake (Table 3).

**Table 3:**
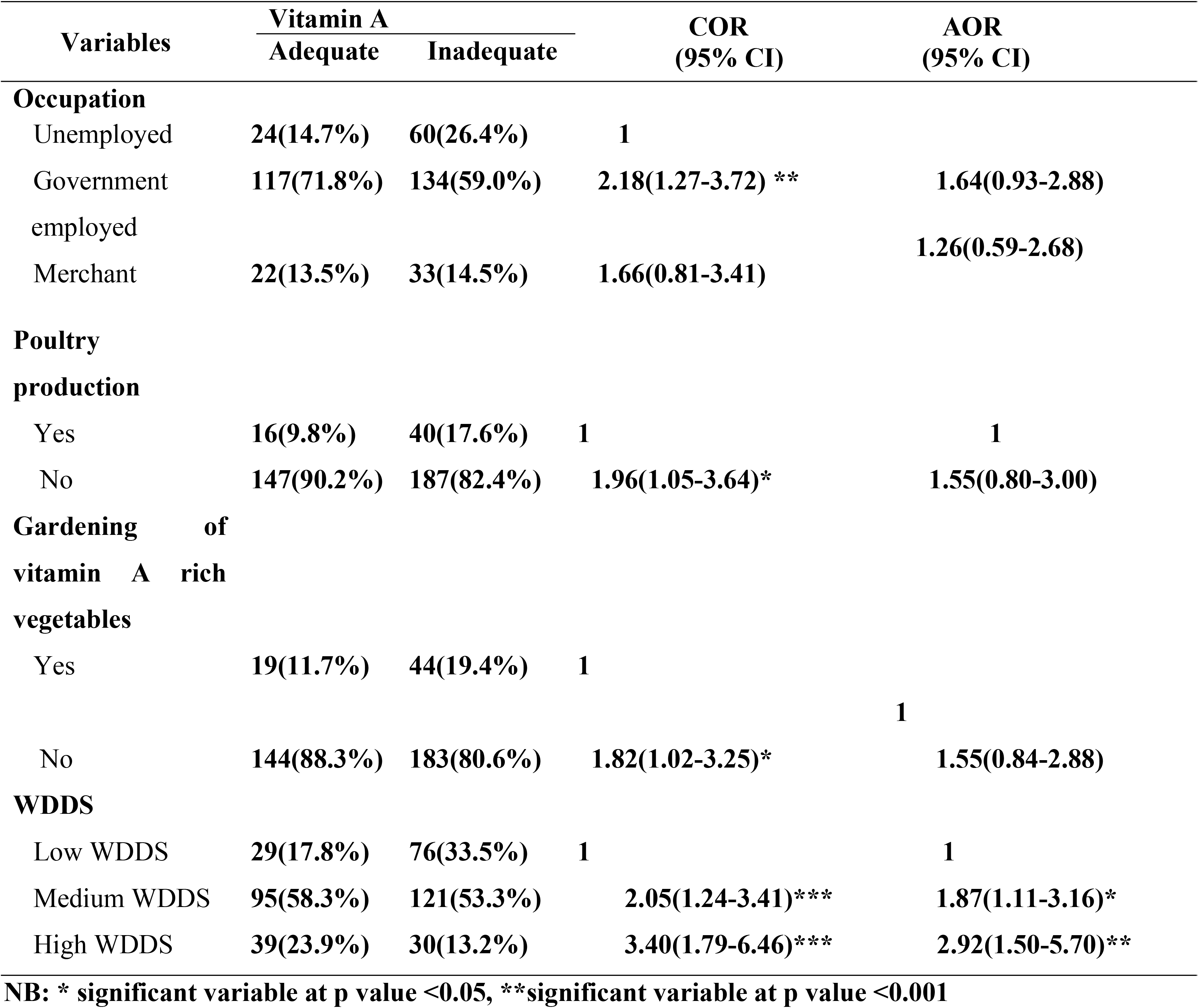
Bivariate and multivariable logistic regression analysis for associated factors of adequacy of vitamin A in pregnant women, health facilities of Dessie town, May 2017.

## Discussions

The current study provides rough estimation of dietary vitamin A intake among pregnant women in which the mean intake was found to be 294.1μg RE/day. This finding was much lower than the mean intake of vitamin A among women in Asian countries like China (19), Korea (20) and Japan (21), European and American countries, such as Italy (22), Spain (23), Mexico (24) and United States (25) that ranged from 480.9 μg RE/day in China to 890 μg RE/day in Italy. The variation among the current finding and findings from the above mentioned countries may due to clear difference in socio-economic status, knowledge and skills for proper dietary practices. The other reason for the discrepancies in proportion of adequate intake of vitamin A may be the use of different recommendations of vitamin A intake in different countries.

Moreover, the percentage nutrient adequacy ratio of vitamin A among pregnant women was about 58%; those pregnant women who cannot meet the mean requirement of vitamin A in μg RE/day of 370 as recommended by joint FAO/WHO consultation (17). This finding was roughly consistent with the findings of the studies conducted in Indonesia central Java (26) and South Africa (27) which were 59.5% and 62.5% respectively. The apparent difference between the two proportions of adequacy may be due to the highest estimated average requirement of South Africa (550mcg RE/day) for pregnant women. The low proportion of adequacy of dietary vitamin A intake were found to be similar low income countries as stated above that may be due the presence of low economy and poor dietary practices and knowledge seeking behavior. However, significant difference in adequacy of vitamin A among pregnant women was observed in Sirilanka (28) as compared to the present study.

The major contributors of vitamin A for consumption among pregnant women were plant sources, particularly vegetables and fruits as pro-vitamin A form and mango took the loin share contribution of Carotenoids. The retinol share of dietary vitamin A was negligible in which the overall consumption of meat, eggs and dairy were below 25%.

In multiple logistic regression analysis, women dietary diversity score was found to be significantly associated with nutrient adequacy ratio of vitamin A. The odds of adequacy of vitamin A among pregnant women with high WDDS was about three times higher than the odds of adequacy of vitamin A among pregnant women with low WDDS. This result is comparable with two studies conducted in Ethiopia, Rural Sidama and Wolaita, Southern Ethiopia (29, 30). The higher diversified diet the women consume, the more nutrient adequacy expected and less risky for micronutrient deficiencies. The result was also in line with the theoretical knowledge that low dietary diversity is related with inadequacy of nutrients including vitamin A. Nutrition specific interventions and nutrition sensitive intervention programs determine the immediate determinants of fetal and maternal health i.e. adequate food and nutrient intake through dietary diversification and modification integrated with sound promotion work.

Even though it was not significant, pregnant women who consume poultry from their produce had the higher dietary intake of vitamin A. consumption poultry directly contribute to the total pool of retinol equivalent without conversion factor (31).

Assessment of vitamin A adequacy measures risk of vitamin A deficiency but not vitamin A status or vitamin A deficiency. How much of beta carotene was absorbed and also converted to RE was not estimated in this study. Left over foods was not estimated. Some foods were estimated by raw portion size. Laboratory assessment along with functional test of vitamin A nutriture investigation was recommended to know the actual burden of the vitamin deficiency among this vulnerable population groups.

## Conclusions

In this study adequacy of vitamin was found to be low which demonstrated a risk of vitamin A deficiency among the studied pregnant women. This signal reinforces a new focus on food based approaches and vitamin A supplementation to reduce the risk of vitamin A deficiency among pregnant women. Women dietary diversity score was predictor of vitamin A adequacy in this study. It is proved that dietary diversity during pregnancy was crucial to increase adequacy of vitamin A.

Advocacy and promotion work is needed to facilitate improved dietary intake by pregnant women; to this end preparing an algorithm for nutrition counseling is the mandate of the top level health department in order to create commonness in the message throughout the health facilities in the studied area.

## List of abbreviations

ANC: Antenatal Care
FAO: Food and Agricultural Organization
NAR: Nutrient Adequacy Ratio
RE: Retinol Equivalent
SD: Standard Deviation
VAD: Vitamin A Deficiency
WDDS: Women Dietary Diversity Score

## Declaration

### Ethics approval and consent to participate

Ethical clearance was obtained from the ethical review committee of the College of Medicine and Health Sciences of Wollo University. All study participants were reassured that names were not needed and would not be recorded. Chances were given to ask anything about the study and been made free to refuse or stop the interview at any moment they want. Informed written consents were obtained from the study participants.

### Consent to Publication

not applicable.

### Availability of Data and Material

the datasets during and/or analyzed during the current study is available from the corresponding author on reasonable request

### Authors’ contribution

ZK: write up the proposal, did the data collection and analysis and involved in manuscript preparation and revision. TCM: involved in the proposal development, data analysis and made the manuscript ready for submission. GE: involved in proposal development and revision of the manuscript. All authors have read and approved the final manuscript’ and All authors contributed equally.

## Acknowledgement

Our special thanks go to data collectors and supervisors. We also acknowledge study participants and health managers at all level of health facilities.

## Acknowledgement

Wollo University was the sponsor of the research. Data collectors, supervisors and health facility managers had played crucial role for the well accomplishment of the research field work.

